# Mapping and targeting of *C1ql1*-expressing cells in the mouse

**DOI:** 10.1101/2023.07.17.549329

**Authors:** Shayan Moghimyfiroozabad, Maëla A. Paul, Séverine M. Sigoillot, Fekrije Selimi

**Affiliations:** Center for Interdisciplinary Research in Biology (CIRB), College de France, CNRS, INSERM, Université PSL, Paris, France

**Keywords:** Mouse model, Cre recombinase, C1ql1, brain, olivo-cerebellar network, rAAV

## Abstract

The C1Q complement protein C1QL1 is highly conserved in mammals where it is expressed in various tissues including the brain. This secreted protein interacts with Brain-specific Angiogenesis Inhibitor 3, BAI3/ADGRB3, and controls synapse formation and maintenance. *C1ql1* is expressed in the inferior olivary neurons that send projections to cerebellar Purkinje cells, but its expression in the rest of the brain is less documented. To map *C1ql1* expression and enable the specific targeting of *C1ql1*-expressing cells, we characterized a knockin mouse model expressing the Cre recombinase under the control of *C1ql1* regulatory sequences. We characterized the capacity for Cre-driven recombination in the brain and mapped Cre expression in various neuron types using reporter mouse lines. Using an intersectional strategy with viral particle injections, we show that this mouse line can be used to target specific afferents of Purkinje cells. As *C1ql1* is also expressed in other regions of the brain, as well as in other tissues such as adrenal glands, placenta, colon and testis, our mouse model is a useful tool to target *C1ql1*-expressing cells in a broad variety of tissues.

## Introduction

C1QL1 belongs to the subfamily of Complement component 1 Q subcomponent-Like proteins (C1QLs). This subfamily is part of the C1Q complement family characterized by a C-terminal globular gC1Q signature domain involved in the formation of hetero-or homo-trimers (1,2). The N-terminal region of C1QL proteins contains two conserved cysteines allowing trimers to form higher-order oligomers (3). C1QLs are absent in plants, yeasts and several invertebrates (e.g. nematodes or insects), while they are found in fish, frog and mammals (3). Among mammals, the C1QL proteins are highly conserved with more than 90% amino acid identity between human and mouse orthologs (3).

In rodents, *C1ql1* is expressed in the brain (3,4), as well as the adrenal glands, placenta, colon and testis (Mouse ENCODE transcription data: https://www.ncbi.nlm.nih.gov/gene/23829). In the central nervous system, its expression is mostly associated with motor-related areas. Some nuclei in the medulla, pons, midbrain, olfactory bulb, hippocampus and cortex express *C1ql1* during postnatal development and adulthood (3,4). In the olivo-cerebellar network, transient expression of *C1ql1* is observed at P7 in the external granular layer (EGL) of the cerebellum, where the precursors of the granule cells reside (3). *C1ql1* is also expressed in inferior olivary neurons (IONs) starting from birth and throughout the life of the mouse (3,4). C1QL1 is a secreted synaptic protein known to be released by the IONs and to bind to its postsynaptic receptor, the Adhesion-G Protein Coupled Receptor (GPCR) Brain-specific Angiogenesis Inhibitor 3 (BAI3, ADGRB3), on cerebellar Purkinje cells (PCs) (4,5). The interaction between C1QL1 and BAI3 is essential for the formation and maintenance of excitatory synapses between climbing fibers (CFs), axons from IONs, and their PC targets, as the removal of each of these two components results in the loss of a significant amount of these synapses (4,5).

In order to fully understand the role of C1QL1 in different brain circuits, it is imperative to be able to specifically target *C1ql1*-expressing cells. To this aim, we developed the *C1ql1*^*Cre*^ knockin mouse model. We characterized the specificity and capacity of this mouse line as a driver using several reporter mouse lines. Finally, we show its usefulness for specific targeting of IONs *in vivo* using an intersectional strategy with viral vectors.

## Results

### Characterization of the driver capacity of the *C1ql1*^*Cre*^ mouse line

To develop a mouse line expressing the Cre recombinase under the control of the *C1ql1* regulatory elements without modifying the coding sequence (CDS) of the *C1ql1* gene, we targeted its last exon (in collaboration with genOway). A transgene was designed to contain two flippase recognition target (FRT) flanking a neomycin (neo) coding sequence, followed by the Exon 2 of the *C1ql1* gene where an IRES-Cre CDS cassette is inserted after the *C1ql1* stop codon. This transgene was knocked in the corresponding region on chromosome 11 by homologous recombination (Fig 1A). The neomycin sequence was removed later by breeding the transgenic mice with a flippase (Flp)-expressing mouse (Fig 1A). Identification of the *C1ql1*^*Cre*^ F0 transgenic mice containing the IRES-Cre CDS was subsequently performed by PCR (Fig 1B). Mice that are homozygous for the *C1ql1*^*Cre*^ alleles are viable, fertile, normal in size and do not show any gross physical or behavioral deficits (observations on two different litters of five animals each, at birth, weaning and adult stages).

**Fig 1.**
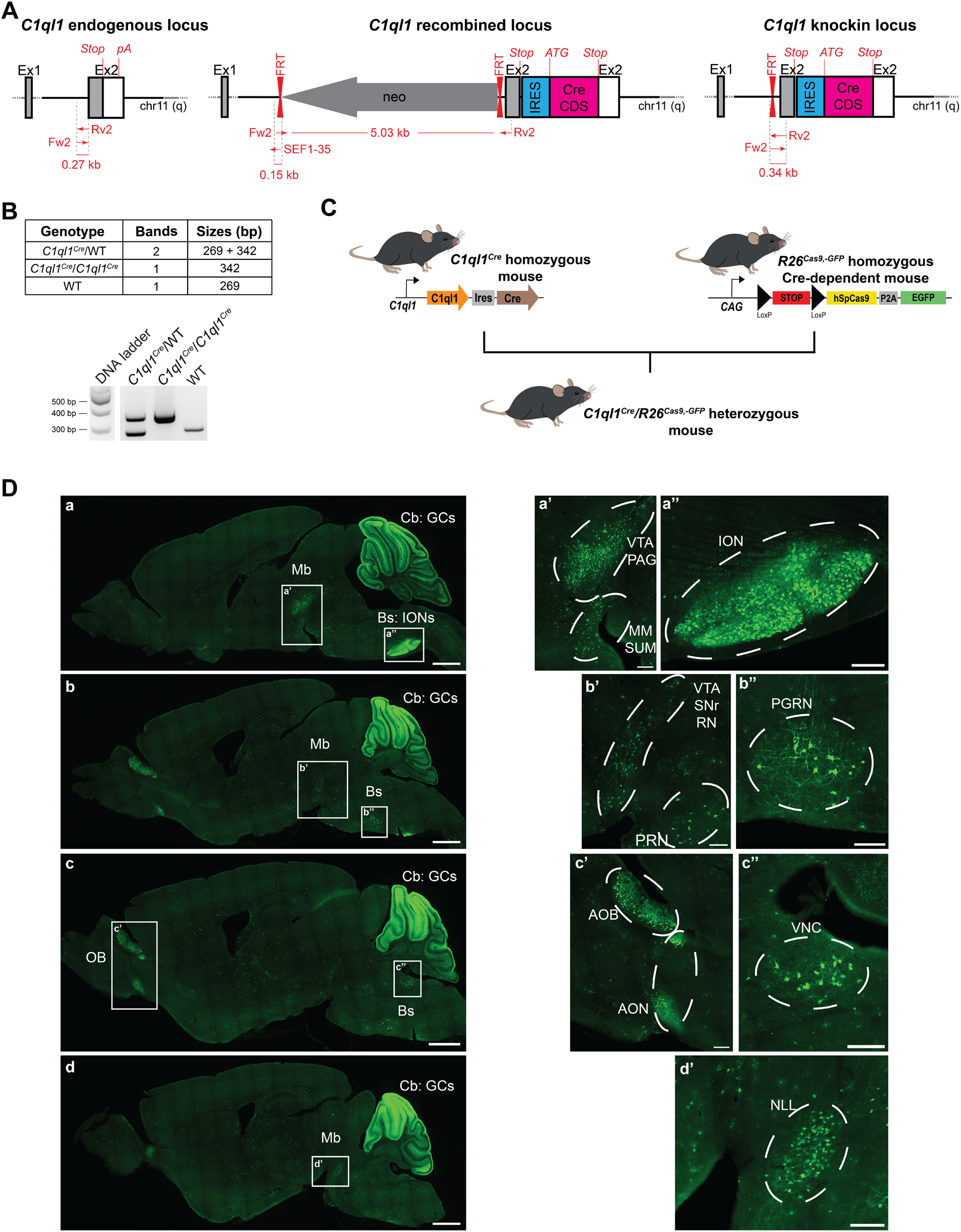
Characterization of the driver capacity of the *C1ql1*^*Cre*^ mouse line. **(A)** Illustration of the mouse *C1ql1* locus at the 2^nd^ exon and flanking regions. Homologous recombination was used to insert a neomycin cassette upstream of the *C1ql1* Exon2 and an IRES-Cre cassette downstream of the stop codon in the *C1ql1* Exon2. After neomycin selection and mice generation, the neomycin cassette was removed using flippase to obtain C1ql1-IRES-Cre knockin mice (*C1ql1*^*Cre*^). **(B)** *Top panel:* Table of the genotyping PCR amplicon bands and their respective sizes depending on the genotype. *Bottom panel:* Representative image of a genotyping PCR run. **(C)** Schematic illustration of the breeding strategy used to generate *C1ql1*^*Cre*^/*R26*^*Cas9,-GFP*^ double heterozygous mice. **(D)** Medial to lateral sagittal sections from P30 *C1ql1*^*Cre*^/*R26*^*Cas9,-GFP*^ mouse brain were immunostained for GFP at P30. Mosaic images were acquired using a Zeiss Axiozoom V16 macroscope (a – d). EGFP was highly expressed in the cerebellum (Cb), in particular in Granule Cells (GCs), and in the Brainstem (Bs), and present in other nuclei from the Midbrain (Mb) and Olfactory Bulb (OB). Scale bars = 1000 μm. (a’ – d’) High magnification of different nuclei or neurons expressing eGFP in P30 *C1ql1*^*Cre*^/*R26*^*Cas9,-GFP*^ mice. Scale bars = 200 μm. AOB: Accessory Olfactory Bulb, AON: Anterior Olfactory Nucleus, ION: Inferior Olivary Nuclei, MM: Medial Mammillary nucleus, NLL: Nucleus of the Lateral Lemniscus, PAG: Periaqueductal Gray, PGRN: Paragigantocellular Reticular Nucleus, PRN: Pontine Reticular Nucleus, RN: Red Nucleus, SNr: Substantia Nigra pars reticulata, SUM: Supramammillary nucleus, VNC: Vestibular Nuclei, VTA: Ventral Tegmental Area.

To visualize and characterize the driver capacity of the *C1ql1*^*Cre*^ mouse line in the brain, homozygous *C1ql1*^*Cre/Cre*^ mice were crossed with the Cre-dependent reporter mouse line *R26*^*Cas9,-GFP/Cas9,-GFP*^ (6) (Fig 1C). In this reporter mouse, the presence of the CRE cleaves the STOP codon before the Cas9-P2A-GFP cassette. Due to the ubiquitous activity of the *Rosa26* locus (*R26*) and the use of the CAG promoter, Cas9 and soluble GFP are expressed permanently in all the cells that have expressed Cre and their progeny. This expression history was mapped in *C1ql1*^*Cre/WT*^/*R26*^*Cas9,-GFP/WT*^ heterozygotes at P30 using immunofluorescence on brain sagittal sections with an anti-GFP antibody to increase the signal to noise ratio (Fig 1D). The GFP-positive cells were observed in different regions of the forebrain, midbrain, brainstem and cerebellum (Fig 1D). At low magnification, two GFP-expressing regions were identified distinctly: the brainstem and the cerebellum (Fig 1D, a). In the brainstem, all the inferior olivary neurons (IONs) expressed GFP strongly (Fig 1D, a), in concordance with previous results showing expression of *C1ql1* amongst these neurons from early development until adult age (3). In the cerebellum, many granule cells (GCs) expressed GFP in a heterogeneous manner and in an antero-posterior gradient, with highest GFP signal in the anterior lobules. Based on *in situ* hybridization data (3), only cells in the external granular layer, not the mature granule cells, express *C1ql1* during the first two postnatal weeks. This suggests that the Cre-dependent recombination happens at the progenitor level in the granule cell lineage. Altogether, these results show that the *C1ql1*^*Cre*^ mouse line is a powerful tool to genetically target two components of the olivo-cerebellar circuit, the IONs and the GCs.

At higher magnification, other GFP-expressing regions were identified in the brain of *C1ql1*^*Cre/WT*^/*R26*^*Cas9,-GFP/WT*^ mice (Fig 1D, a’-d’). According to the Allen Reference Brain atlas (https://mouse.brain-map.org/static/atlas), we found GFP-expressing cells in several motor-related areas in the midbrain: ventral tegmental area, periaqueductal gray (Fig 1D, a and a’), substantia nigra pars reticulata and red nucleus (Fig 1D, b and b’). Adjacent to these regions, several GFP-positive cells were detected in two medial nuclei of the hypothalamus: supramammillary nucleus and medial mammillary nucleus (Fig 1D, a and a’). In the pons, two nuclei were labelled with GFP: pontine reticular nucleus (motor related-area) (Fig 1D, b and b’) and nucleus of the lateral lemniscus (sensory-related area) (Fig 1D, d and d’). In addition to the IONs (Fig 1D, a and a’’), two other motor-related nuclei were expressing GFP in the medulla: paragigantocellular reticular nucleus (Fig 1D, b and b’’) and vestibular nuclei (Fig 1D, c and c’’). Expression of GFP is not restricted to the midbrain or hindbrain, but can be found also in the forebrain, in the accessory olfactory bulb and anterior olfactory nucleus (Fig 1D, c and c’). The same pattern of expression was found when *C1ql1*^*Cre/Cre*^ mice were crossed with another reporter line (Supp Fig 1), showing the reproducibility of neuronal targeting using the *C1ql1*^*Cre*^ mouse line. Altogether, different nuclei and regions, especially motor-related areas in the midbrain, pons and brainstem, can be targeted genetically with the *C1ql1*^*Cre*^ mouse model, in addition to the olivo-cerebellar network.

### Normal morphology and transmission of climbing fiber/Purkinje cell synapses in *C1ql1*^*Cre/WT*^ heterozygotes

To detect the expression of the Cre recombinase in different brain regions, RT-qPCR was performed on RNA extracts from the cerebellum, the brainstem, the hippocampus and the cortex at P14. No expression of Cre was observed in extracts from control (wild-type, WT) mice, while Cre was detected in all tested regions of the brain in heterozygous *C1ql1*^*Cre/WT*^ animals (Fig 2A). As expected, the highest expression of Cre was seen in the brainstem extracts where IONs are located (mean expression normalized to *Rpl13a* ± SEM = 6.40 ± 0.54 in the brainstem, *versus* 2.54 ± 0.54, 2.60 ± 0.57, and 1.53 ± 0.18 in the cerebellum, the hippocampus, and the cortex, respectively). C1QL1 loss of function leads to detectable phenotypes in CF/PC synapse formation and function in the olivo-cerebellar network when *C1ql1* expression is reduced by more than 90% (4,5). We assessed by RT-qPCR whether the insertion of the transgene had any impact on the expression of the endogenous *C1ql1* gene. At P14 in extracts from heterozygous animals, *C1ql1* mRNA levels were decreased by 50-60% in all regions tested, compared to WT animals (Fig 2A), showing that the insertion of the transgene prevents the proper expression of *C1ql1 m*RNA from the locus where it is inserted. To test whether this level of decrease leads to functional consequences, we performed morphological and functional analysis of climbing fiber/Purkinje cell (CF/PC) synapses comparing *C1ql1*^*WT/WT*^ and *C1ql1*^*Cre/WT*^ heterozygotes. Morphological analyses were performed in two lobules of the cerebellum: lobules VI and IX, at P30 when the olivo-cerebellar circuit is established (7), using immunofluorescence against vesicular glutamate transporter 2 (VGLUT2), to label the presynaptic boutons of CFs, and against calbindin (CaBP) to label PCs (Fig 2B, *left panel*). Quantification showed no major difference in the thickness of molecular layer of the cerebellar cortex and of the extension of CF innervation on the dendritic tree of PCs (Fig 2B, *right panel*). While loss of function of *C1ql1* at a level >90% leads to decreased density of VGLUT2 clusters and decreased transmission (4,5), our quantification did not detect any differences in the volume and density of VGLUT2 labelled presynaptic boutons between *C1ql1*^*WT/WT*^ and *C1ql1*^*Cre/WT*^ heterozygotes (Fig 2B, *right panel*). Patch-clamp recordings of CF/PC transmission during the fourth postnatal week showed similar amplitude of EPSCs in slices from *C1ql1*^*WT/WT*^ and *C1ql1*^*Cre/WT*^ heterozygote mice (Fig 2C). Altogether, the expression of Cre and concomitant decrease of *C1ql1* expression in *C1ql1*^*Cre/WT*^ heterozygous animals do not impact morphological or electrophysiological properties of CF/PC synapses. The *C1ql1*^*Cre*^ mouse line can thus be used to manipulate the olivo-cerebellar circuit when used as heterozygote without detrimental effect on normal development and function of the circuit.

**Fig 2.**
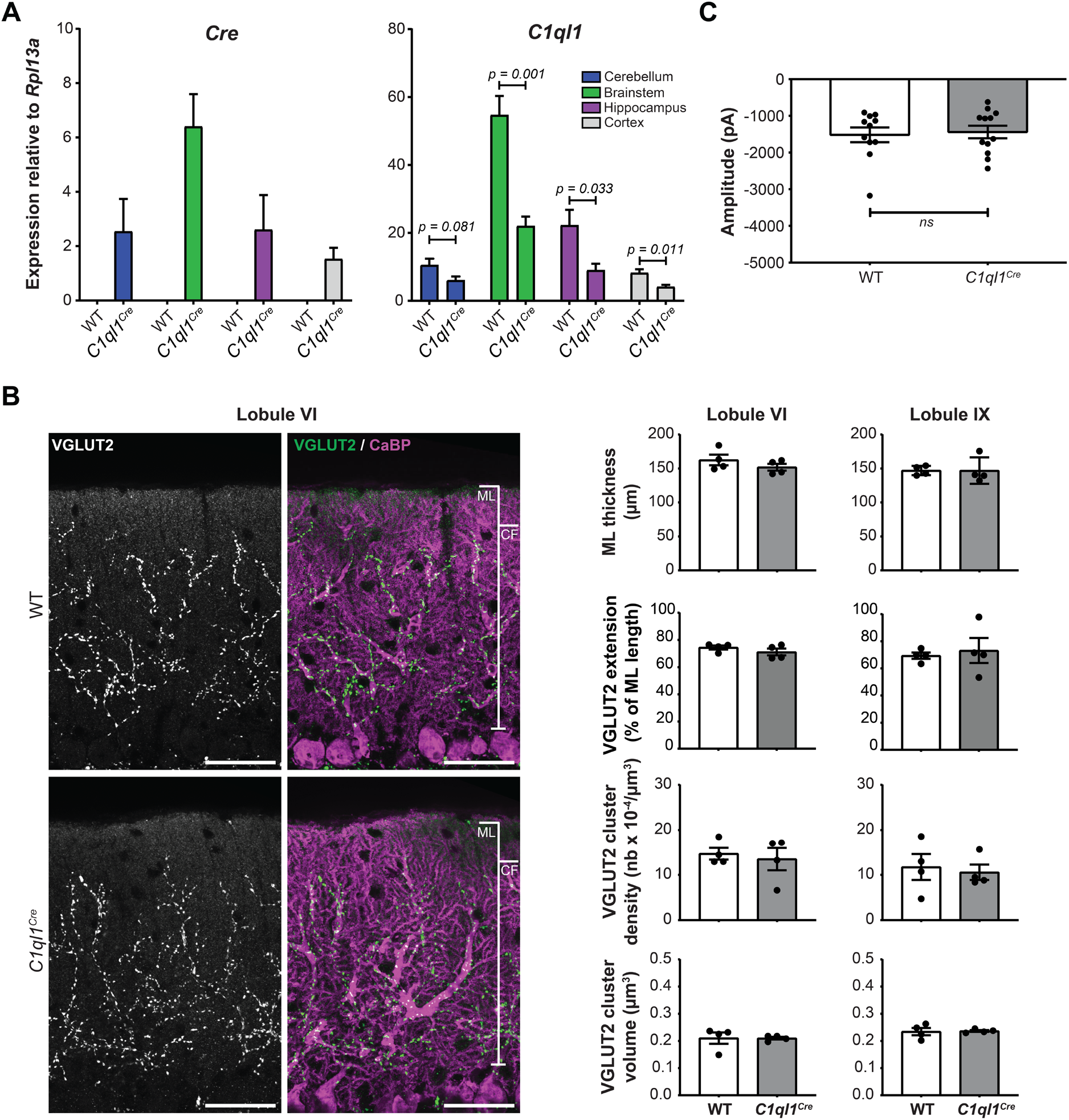
No effect of Cre transgene knockin on the morphology and transmission of climbing fiber/Purkinje cell synapses in *C1ql1*^*Cre*^ heterozygotes. **(A)** The expression of *Cre* and *C1ql1* mRNAs was assessed using quantitative RT-PCR on RNA extracts from four different regions of *C1ql1*^*WT/WT*^ (WT) or *C1ql1*^*Cre/WT*^ (*C1ql1*^*Cre*^) mouse brains, fourteen days after birth (P14). The *Cre* recombinase expression was detected only in *C1ql1*^*Cre*^ mice, in the cerebellum, the brainstem, the hippocampus and the cortex. The expression of *C1ql1* was decreased in these regions of the brain at the same age compared to WT mice. Data are represented as mean ± SEM, unpaired student t test with Welsh’s correction, n = 5 – 6 animals per genotype. **(B)** *Left panel:* CF presynaptic boutons and PCs and their dendritic tree were revealed by immunostaining for VGLUT2 (green) and calbindin (magenta) respectively, in parasagittal cerebellar sections from P30 WT and *C1ql1*^*Cre*^ mice. Images were acquired using a spinning disk confocal microscope. *Right panel:* analyses of the morphology of the cerebellar cortex and connectivity of CFs were performed on cerebellar lobules VI and IX. The thickness of molecular layer, territory of the CF synapses on PC dendritic tree, volume and density of the boutons of CFs on PCs were not modified in lobules VI and IX of *C1ql1*^*Cre*^ heterozygous animals compared to WT. Molecular layer (ML) and CF synapse extension (CF) are indicated on the merge images. Data are represented as mean ± SEM, unpaired student t test with Welsh’s correction, n = 4 animals per genotype. Scale bars: 40 µm. **(C)** Properties of CF/PC transmission was assessed in WT and heterozygous *C1ql1*^*Cre*^ mice recording the mean amplitude of the currents of CF on PCs. This analysis revealed no difference between the two genotypes. Data are represented as mean ± SEM, Mann Whitney test, n = 11 – 12 cells per genotype.

### Targeting different cell types in the olivo-cerebellar network using the *C1ql1*^*Cre*^ mouse line

The *C1ql1*^*Cre*^ mouse line is expected to drive the expression of the Cre recombinase in the olivo-cerebellar network, and in particular in IONs and GC precursors. To confirm this, we crossed *C1ql1*^*Cre/Cre*^ homozygous mice with *R26*^*hM4Di,-mCitrine/WT*^ (8) or *R26*^*R-EYFP/WT*^ (9) reporter mice (Fig 3). In *C1ql1*^*Cre/WT*^/*R26*^*hM4Di,-mCitrine/WT*^ heterozygous mice, the DREADD hM4Di and soluble mCitrine are co-expressed in cells following the excision by the Cre of the Stop cassette preceding the hM4Di-P2A-mCitrine sequence (8). Immunofluorescence on brain sagittal sections at P30 using an anti-GFP antibody revealed expression of mCitrine in the inferior olive in the brainstem, with clear labeling of CFs in this region (Fig 3A). mCitrine-positive GCs were also observed in the internal granular layer of the cerebellum (Fig 3A). In *C1ql1*^*Cre/WT*^/*R26*^*R-EYFP/WT*^ heterozygous mice, the enhanced YFP is expressed in cells following excision by the Cre of the Stop cassette preceding the EYFP sequence (9) and immunofluorescence experiments showed similar expression of EYFP in IONs and CFs, as well as in cerebellar GCs (Fig 3B). Of note, we observed in some instances mCitrine expression in the Bergmann glia in the cerebellum when breeding a *R26*^*hM4Di,-mCitrine/WT*^ male with *C1ql1*^*Cre/Cre*^ females but not with the reverse crossing (data not shown). This type of parental sex-related differences was not observed when breeding to the *R26*^*Cas9,-GFP*^ reporter line, suggesting differences linked to the reporter mouse lines.

**Fig 3.**
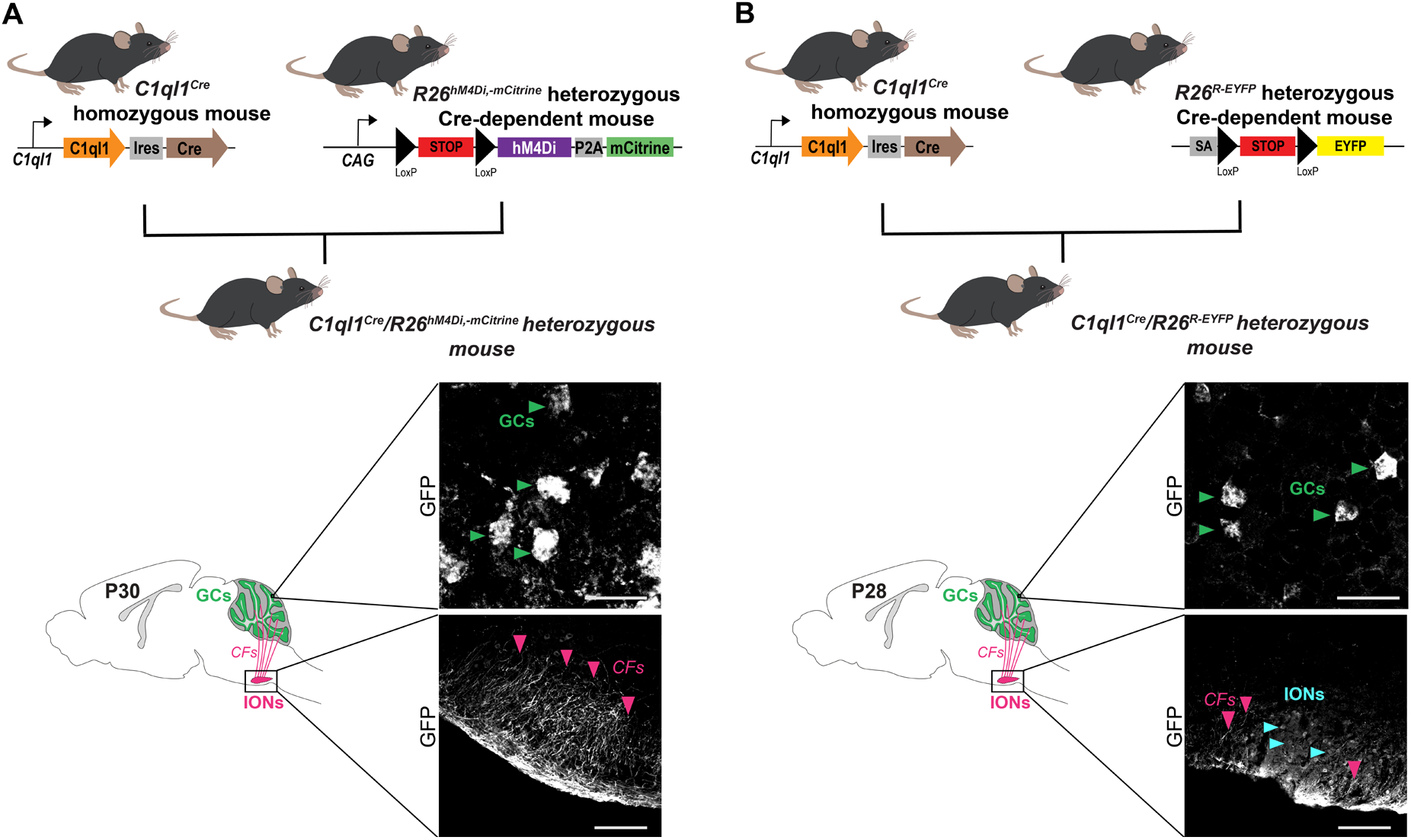
*C1ql1*^*Cre*^ mouse line drives expression in different cell types of the olivo-cerebellar network. **(A)** *Top panel:* Schematic illustration of the crossings used to generate *C1ql1*^*Cre*^/*R26*^*hM4Di,-mCitrine*^ heterozygous mice. *Bottom panel:* parasagittal brain sections from P30 *C1ql1*^*Cre*^/*R26*^*hM4Di,- mCitrine*^ mice were immunostained for mCitrine using anti-GFP antibody and images were acquired using a spinning disk confocal microscope. Cerebellar granule cells (GCs, green arrowheads) and axons from inferior olivary neurons (IONs), the climbing fibers (CFs, pink arrowheads), were positive for mCitrine in the cerebellum and brainstem regions, respectively. **(B)** *Top panel:* Schematic illustration of the crossings used to generate *C1ql1*^*Cre*^/*R26*^*R-EYFP*^ heterozygous mice. SA, Splice Acceptor. *Bottom panel:* parasagittal cerebellar sections from P28 *C1ql1*^*Cre*^*/R26*^*R-EYFP*^ mice were immunostained for EYFP using anti-GFP antibody. Cerebellar GCs (green arrowheads) and CFs (pink arrowheads) from IONs (cyan arrowheads) were EYFP labelled in the cerebellum and brainstem regions, respectively. Scale bars = 20µm for GCs and 125 µm for IONs.

To specifically target the IONs using *C1ql1*^*Cre/WT*^ heterozygous mice, we developed an intersectional strategy. A retrograde AAV driving the expression of the Cre-dependent VAMP2-GFP construct under the CamKII promoter was injected in the cerebellum of postnatal day 0 (P0) mouse pups or P27 juvenile mice. As CamKII is expressed in IONs but not in GCs, the conditional expression of VAMP2-GFP was specific to the inferior olive, both at P27 and P44 after P0 and P27 injections, respectively (Fig. 4A). While the breeding of *C1ql1*^*Cre*^ with, for example, the reporter mouse line *R26*^*Cas9,-GFP*^ led to labelling of the entire ION population (Fig 1D, a and a’’), only subpopulations of IONs are targeted using the intersectional strategy. Using immunofluorescent labeling of brain sagittal sections at P27 or P44 (data not shown), with an anti-GFP antibody (Fig 4B), GFP-positive climbing fibers were observed in different lobules in the cerebellum, in particular in lobules V to X in the vermis or paravermis (Fig 4B, *top panel*). At higher magnification, GFP-positive climbing fibers were visualized along the proximal dendrites of PCs (Fig. 4B, *bottom panel*). No GFP-positive GCs were observed, showing the high specificity of expression in one of the two excitatory afferents of the PCs. Altogether, our results show that the *C1ql1*^*Cre*^ mouse line can be used to either target both PC excitatory inputs or only one of them, depending on the strategy that is used.

**Fig 4.**
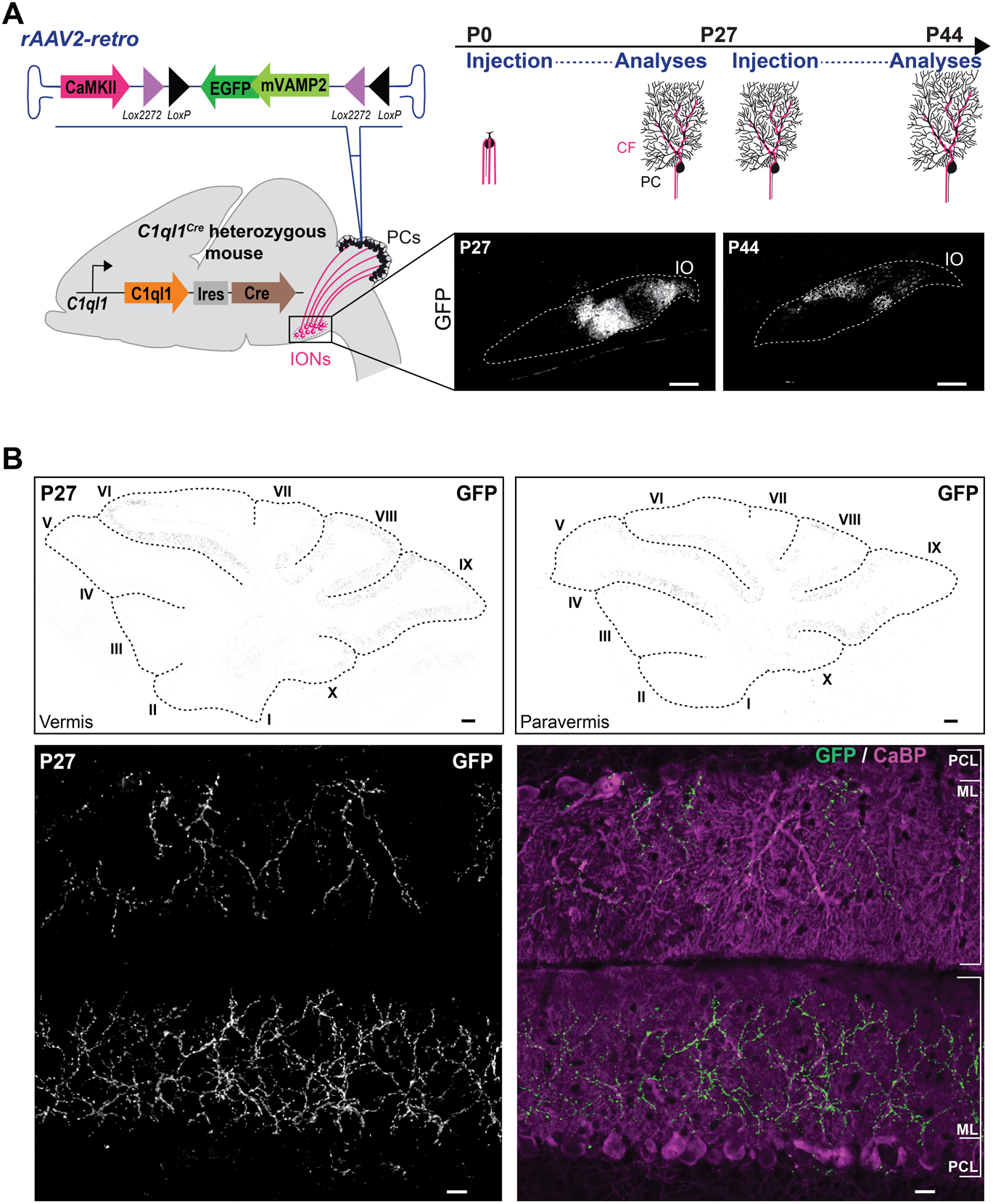
Specific targeting of inferior olivary neurons using an intersectional strategy. **(A)** Schematic illustration of the strategy to specifically express the mouse GFP-tagged vesicle-associated membrane protein 2 (mVAMP2) in IONs. P0 or P27 stereotaxic injections of retrograde rAAVs expressing the floxed mVAMP2-GFP, under the CaMKII promoter, were performed in the cerebellum of *C1ql1*^*Cre*^ mice. Analyses were done at P27 for P0 injections or P44 for P27 injections, in parasagittal cerebellar sections. Subpopulations of IONs were labelled for GFP after anti-GFP immunostaining. Inferior Olive (IO) is indicated by dotted lines. Scale bars, 200 µm. **(B)** *Top panel:* Vermal (left) and paravermal (right) sagittal cerebellar sections from *C1ql1*^*Cre/WT*^ P27 mouse injected at P0 and immunostained for GFP revealed large region of GFP labelled CFs in different lobules of the cerebellum. Images were acquired using a spinning disk confocal microscope. Lobules are indicated as roman numerals. Scale bars, 500 µm. *Bottom panel:* CFs of transduced IONs and Purkinje cells from P27 *C1ql1*^*Cre/WT*^ mouse injected at P0 were immunostained for GFP (green) and calbindin (magenta), respectively. Molecular Layer (ML) and Purkinje cell layer (PCL) are indicated on the merge image. Scale bars = 20 µm.

## Discussion

C1QL1 is a secreted protein involved in synapse formation and maintenance between inferior olivary neurons and Purkinje cells in the olivo-cerebellar circuit (4,5). Since the expression of *C1ql1* was identified in inferior olivary neurons and in the external granular layer during development, we developed a *C1ql1*^*Cre*^ mouse model by inserting an IRES-Cre cassette after the stop codon of *C1ql1*, to enable the genetic manipulation of the two sources of excitatory inputs to PCs, IONs and GCs, *in vivo*. Using RT-qPCR, the expression of Cre is detected in the cerebellum, brainstem, hippocampus and cortex. The specificity and capacity of the *C1ql1* driver was characterized by crossing the *C1ql1*^*Cre*^ mouse line with three CRE-dependent reporter mouse models: *R26*^*Cas9,-GFP*^, *R26*^*hM4Di,-mCitrine*^ and *R26*^*R-EYFP*^. The expression of the reporter proteins is observed not only in IONs and GCs, but also in some other nuclei in the brainstem, midbrain and olfactory bulb. To specifically target IONs during postnatal development and adulthood, a viral strategy has been developed *via* stereotaxic injection of a retrograde rAAV2 with a CRE-dependent reporter gene in the cortex of the cerebellum at P0 or P27, resulting in the transduction of numerous IONs and CFs 2-4 weeks after the injection. In general, the *C1ql1*^*Cre*^ mouse model is a powerful genetic tool to target and manipulate *C1ql1*-expressing cells in the brain, in particular IONs in the brainstem.

Using the CRE-dependent reporter mouse models to map the location of *C1ql1*-expressing cells, we show the localization of the reporter gene in several nuclei in different regions of the brain. These nuclei are involved in different functions (e.g. olfaction, arousal and consciousness), but some of them found in the brainstem, cerebellum, pons and midbrain, such as inferior olive, red nucleus, substantia nigra pars reticulata and GCs, are motor-related structures. We have not analyzed the expression of *C1ql1* and *Cre* in other organs than the brain in *C1ql1*^*Cre*^ mouse model, but several studies show the expression of *C1ql1* in testis, liver (3), adipose tissue (10), the adrenal glands, colon (https://www.ncbi.nlm.nih.gov/gene/23829), and ovaries (11). The function of C1QL1 in ovarian folliculogenesis has been investigated in mice using a full knockout model, showing its anti-apoptotic effect on granulosa cell apoptosis (11). Loss of function of *C1ql1* using a *C1ql1* constitutive knockout mouse model results in abnormal hair cell innervation in the cochlea and loss of audition (12). In addition to physiological conditions, several studies suggest that C1QL1 might be involved in some malignancies. C1QL1 is upregulated in colorectal cancer, one of the most common malignant cancers (13), differentiated thyroid carcinoma (14), as well as lung adenocarcinoma (15). RNA-seq analyses have shown that *C1ql1* expression is higher in glioblastoma compared to the normal brain (13). All these studies show the diversity in the function of C1QL1 in health and disease in different systems of the body. Thus, *C1ql1*^*Cre*^ mouse is a potentially useful model to study the role of *C1ql1*-expressing cells in different physiological and pathological conditions.

## Methods and Materials

### Animals

All mice were kept in the authorized animal facilities of CIRB, College de France, under a 12 hours light: 12 hours dark cycle with water and food supplied ad libitum. All animal protocols and animal facilities were approved by the Comité Régional d’Ethique en Expérimentation Animale (#2001) and the veterinary services (D-75-05-12). *C1ql1*^*Cre*^ mouse model was generated and provided by genOway. The Cre-dependent reporter mouse lines, *R26*^*Cas9-GFP*^ (B6J.129(B6N)-Gt (ROSA)26Sor^tm1(CAG-cas9*,-EGFP)Fezh^/J, strain #026175) (6), *R26*^*hM4Di,-mCitrine*^ (B6.129-Gt(ROSA)26Sor^tm1(CAG-CHRM4*,-mCitrine)Ute^/J, strain #026219) (8) and *R26*^*R-EYFP*^ (B6.129×1-Gt(ROSA)26Sor^tm1(EYFP)Cos^/J, strain #:006148) (9) were obtained from The Jackson Laboratory. All the lines are maintained on the C57BL/6J background. In all experiments, littermate mice from both sexes were used as controls for the analysis.

### Viral particles and Stereotaxic injections

Injections of AAV particles were performed in the cerebellar vermis of ice-anesthetized P0 *C1ql1*^*Cre/WT*^ heterozygous mice or isoflurane-anesthetized P27 *C1ql1*^*Cre/WT*^ heterozygous mice, at 1 mm depth from the skull and 3.2 mm relative to Bregma or at 1.5 mm depth from drilled skull and 2.7 mm from Lambda respectively, to target Purkinje cell layer. AAV2/retrograde serotype was used to target IONs. 0.25µl of AAV was injected per animal using pulled calibrated pipets. AAVrg-CamKII-DIO-mVAMP2/eGFP-WPRE expressing the floxed mVAMP2-GFP, under the CaMKII promoter were obtained from Vector Biolabs: # AAV-275950 (RefSeq # BC055105), with a titer of 3.2×10^13 GC/mL.

### Generation of C1ql1-IRES-Cre (C1ql1^Cre^) knockin mouse

*C1ql1*^*Cre*^ mice were generated using homologous recombination (genOway). A targeting vector containing Neomycin positive-selection cassette flanked with two FRT sites, and Exon2 of *C1ql1* gene where an IRES-Cre cassette was inserted downstream of the stop codon in this exon, was electroporated into ES cells. The homologous recombination in the targeted ES cell clones was validated by PCR and DNA sequencing. The recombined ES cells with the correct sequence were injected into blastocysts. These blastocysts were then implanted in pseudo-pregnant females. Chimerism rate was then assessed in the progeny by coat color markers comparison. Highly chimeric males (chimerism rate above 50%) were generated and crossed with C57BL/6J Flp deleter female mice were used to excise the Neomycin selection cassette and to generate heterozygous mice carrying the Neo-excised Neo-devoid Knockin. 16 heterozygous mice were screened by PCR followed by sequencing to make sure of the germline transmission.

PCR genotyping is carried out by PCR using two primers (forward primer: 5’-GCCCAGATGTATTCTGCCCTAGAATCC-3’; reverse primer: 5’-ATTGCACTGGCCCGCACCTAAG-3’) to detect the WT (269 bp) and the targeted knock-in alleles (342 bp) (Fig. 1B). Transgenic founders were backcrossed and maintained on the C57Bl/6J background.

### Characterization of C1ql1^Cre^ knockin mouse

Gene expression analysis

#### RNA extraction and cDNA synthesis

The cerebellum, brainstem, hippocampus and a piece of cortex were extracted from *C1ql1*^*WT/WT*^ or *C1ql1*^*Cre/WT*^ P14 mice and frozen immediately in liquid nitrogen, and stored at -80C°. Using the Qiagen RNeasy mini kit (Qiagen, Venlo, Netherlands, #74104), the total RNA was purified according to manufacturer’s instruction. cDNA was synthetized using 100 ng of total RNA and SuperScript™ VILO™ cDNA Synthesis Kit (Life Technologies, California, USA, #11754050) according to manufacturer’s instruction.

#### qPCR

Quantitative PCR on cDNA samples was performed using FAM-labeled *C1ql1* (Taqman Gene Expression Assay, ThermoFisher, Assay ID: Mm00657289_m1) or *Cre* (Taqman Gene Expression Assay, ThermoFisher, Assay ID: Mr00635245_cn) probes and VIC-labeled *Rpl13a* probe as a reference (Taqman Gene Expression Assay, ThermoFisher, Assay ID: Mm01612986-gH) and the TaqMan Universal Master Mix II, with UNG (Applied Biosystems, #4440038) according to manufacturer’s instructions. Bio-Rad CFX Manager was used for data analysis. The relative expression of *C1ql1* to *Rpl13a* or *Cre* to *Rpl13a* was calculated based on the following formula: Relative expression = (2^(Cq of *C1ql1 or Cre* – Cq of *Rpl13a)*^) x 100.

#### Immunofluorescence

Mice tissues were fixed using intracardiac perfusion of 4% PFA in PBS solution. Brains were extracted, post-fixed with the same solution at 4°C for 2 -4 hours, and transferred to 30% sucrose/PBS for 48 hours at 4°C for cryoprotection. 30 µm-thick sections were obtained using a freezing microtome, and kept in 0.02% NaN_3_ in PBS solution at 4°C until use.

To perform immunolabeling, sections were incubated in blocking buffer (4% donkey serum and 1% Triton X-100 in PBS solution) for 30 min to 1 hour at room temperature, followed by incubation with primary antibodies in 1% donkey serum and 1% Triton X-100 in PBS for overnight at 4°C with agitation. Primary antibodies were: GFP (1:1000, chicken, ab13970, Sigma or 1:2000, rabbit, ab6556, Sigma), CaBP (1:2000, rabbit, CB38, Swant or 1:2000, mouse, #300) and VGLUT2 (1:7000, guinea pig, AB2251, Millipore). The slices were washed 3 times in 1% Triton X-100 in PBS for 5 to 10 minutes, followed by incubation with secondary antibodies (Alexa Fluor 488-, and 568- and 647-labeled donkey/goat anti-mouse, rat, rabbit, or chicken IgGs (H+L); 1:1000, Invitrogen or Life Technologies) in 1% Triton X-100 in PBS for 1 to 2 hours at room temperature. Then, sections were washed 3 times in 1% Triton X-100 in PBS for 5 to 10 minutes, and incubated for another 10 min at room temperature with Hoechst 33342 (0,2 mg/mL, Sigma, Gothenburg, Sweden, cat#H6024) and 0.4% Triton X-100 in PBS. Sections were mounted using ProLong Gold Antifade Reagent (Invitrogen, cat#P36930).

#### Image acquisition and analysis

The mosaic images for global brain morphology (Fig 1D and Supp Fig 1) were obtained using a Zeiss Axiozoom V16 macroscope, equipped with a digital camera (AxioCam HRm) using a 160x (pixel size: 0.4037 μm), and reconstructed using the Zeiss Zen software. Images for VGLUT2 quantifications (Fig 2B) were acquired using a Zeiss spinning-disk confocal CSU-W1 microscope with 63x oil objective (Z-plane step size: 0.19 μm). Images for GFP analyses were acquired using the same spinning-disk confocal microscope with 63x or 25x oil objectives (single plane, Fig 3 and Fig 4). The mosaic images for GFP visualization were reconstructed using the stitching program from metamorph software (Fig. 4B *top panel*).

VGLUT2 quantification (cluster density and volume) were performed as described previously (16). All the images were normalized using the quantile-based normalization plugin in Fiji. The intensity distribution of the images has been normalized using 256 quantiles for each staining. The synaptic boutons were extracted from the background using the 3D Weka Segmentation plugin (https://imagej.net/Trainable_Weka_Segmentation) after manual selection of signal and background samples. The Fiji built-in plugin 3D object counter was then used in order to count and measure every object (cluster of VGLUT2 positive signal). CF territory and ML thickness were measured manually using Fiji. All the steps were performed in blind condition.

### Electrophysiology

Acute parasagittal cerebellar slices were obtained from *C1ql1*^*Cre*^ mice from P25 to P30. 200 μm-thick slices were cut at room temperature with a Campden Ci 7000 smz microtome in (in mM): Sucrose 120, NaCl 60, KCl 2.5, D(+)Glucose 25, NaHCO3 25, NaH2PO4 1.25, CaCl2 0.1, MgCl2 3, ascorbic acid 0.4, myo-inositol 3, NaPyruvate 2, pH=7.3-7.4. Slices were then transferred and allowed to recover for one hour at room temperature in the following solution (in mM): NaCl 125, KCl 2.5, D(+)Glucose 25, NaHCO3 25, NaH2PO4 1.25, CaCl2 2, MgCl2 1, ascorbic acid 0.4, myo-inositol 3, NaPyruvate 2, pH=7.3-7.4, oxygenated. This solution was used to fill the stimulation pipette and, complemented with picrotoxin (100 µM), was used as external solution for 5 recordings. Borosilicate glass pipettes with 2-5 MΩ resistance were used for recordings and filled with the following internal solution (in mM): CsCl2 155, Hepes 10, EDTA 5, QX314 5, pH=7.35 adjusted with CsOH. Responses to CF stimulation were recorded at a holding membrane potential of −10 mV in Purkinje cells of lobule VI using a MultiClamp 700B amplifier (Molecular Devices, CA) and acquired using the freeware WinWCP written by John Dempster (https://pureportal.strath.ac.uk/en/datasets/strathclyde-electrophysiology-software-winwcp-winedr). Series resistance was compensated by 80–90% and cells were discarded if significant changes were detected. CF-mediated responses were identified by the typical all-or-none response and strong depression displayed by the second response elicited during paired-pulse stimulations (20 Hz). Electrophysiological data were analyzed using the software Clampfit 10.7 (Molecular Devices).

### Statistical analysis

All the data were analyzed and represented using GraphPad Prism. They are represented as mean ± SEM. The student’s t-test was used to measure the differences between two groups when they had normal distribution. The non-parametric Mann-Whitney test was used when the two groups didn’t have normal distribution.

## Acknowledgments

We would like to thank Yves Dupraz for the development of tools for stereotaxic injections in neonates, Laurent Venance for sharing *R26*^*R-EYFP*^ mice, and the personnel from the CIRB animal and imaging facilities.

## Funding

This work was supported by funding from:

European Research Council ERC consolidator grant SynID 724601 (FS) Q-life ANR-17-CONV-0005 (FS)

ANR-10-LABX-54 MEMO LIFE (FS) INCA PEDIAHR21-014 (FS)

Sorbonne Université ED158 (MAP) Collège de France (MAP)

Ecole des neurosciences de Paris Ile-de-France (SM) La Ligue Nationale Contre le Cancer (SM)

## Author Contributions

Conceptualization: FS Methodology: FS, SM, SMS, MAP Investigation: SM, SMS, FS, MAP Visualization: SM, SMS, FS Funding acquisition: FS

Project administration: FS Supervision: FS and SMS

Writing – original draft: SM and SMS Writing – review & editing: SMS, SM, FS

## Data and materials availability

All data are available in the main text.

---

**Supp Fig 1.**
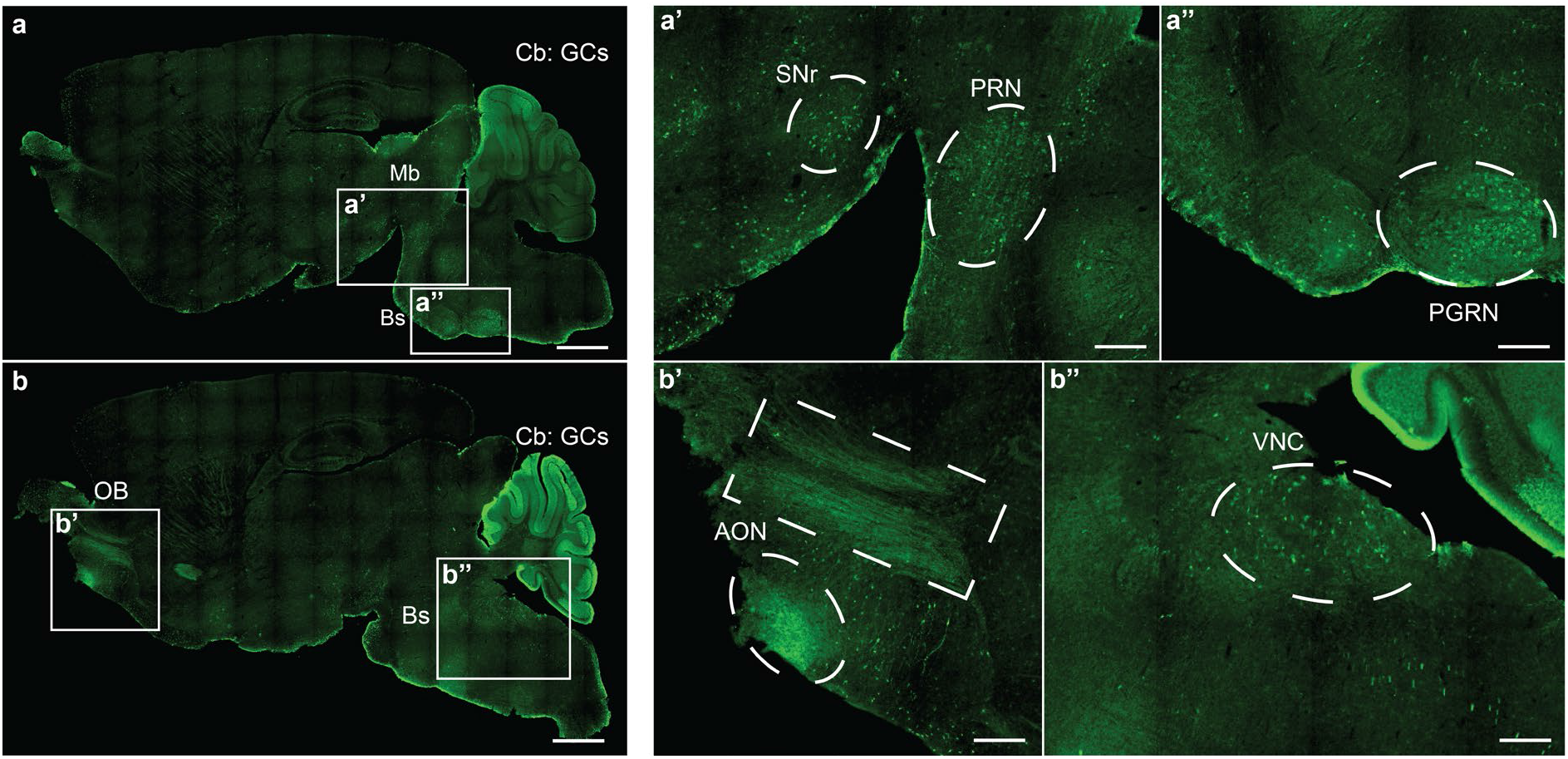
Reproducibility of the targeting capacity of the *C1ql1*^*Cre*^ mouse line. Medial to lateral sagittal sections from P28 *C1ql1*^*Cre*^/*R26*^*R-EYFP*^ mouse brain were immunostained for EYFP using anti-GFP antibody. Mosaic images were acquired using a Zeiss Axiozoom V16 macroscope (a – b). EYFP was highly expressed in the Cerebellum (Cb), in particular in Granule Cells (GCs) and in the Brainstem (Bs), and present in other nuclei from the Midbrain (Mb) and Olfactory Bulb (OB). Scale bars = 1000 μm. (a’ – b’’) High magnification of different nuclei or neurons expressing EYFP in P28 *C1ql1*^*Cre*^/*R26*^*R-EYFP*^ mice. Scale bars = 250 μm. AON: Anterior Olfactory Nucleus, PGRN: Paragigantocellular Reticular Nucleus, PRN: Pontine Reticular Nucleus, SNr: Substantia Nigra pars reticulata, VNC: Vestibular Nuclei.

